# Stimulus-induced changes in 1/*f*-like background activity in EEG

**DOI:** 10.1101/2021.12.17.473188

**Authors:** Máté Gyurkovics, Grace M. Clements, Kathy A. Low, Monica Fabiani, Gabriele Gratton

## Abstract

Research into the nature of 1/*f*-like, non-oscillatory electrophysiological activity has grown exponentially in recent years in cognitive neuroscience. The shape of this activity has been linked to the balance between excitatory and inhibitory neural circuits, which is thought to be important for information processing. However, to date, it is not known whether the presentation of a stimulus induces changes in the parameters of 1/*f* activity, which are separable from the emergence of event-related potentials (ERPs). Here, we analyze event-related broadband changes in scalp-recorded EEG both before and after removing ERPs to demonstrate their confounding effect, and to establish whether there are genuine stimulus-induced changes in 1/*f*. Using data from a passive and an active auditory task (n=23), we found that the shape of the post-event spectra differed significantly from the pre-event spectra even after removing the frequency-content of ERPs. Further, a significant portion of this difference could be accounted for by a rotational shift in 1/*f* activity, manifesting as an increase in low and a decrease in high frequencies. Importantly, the magnitude of this rotational shift was related to the attentional demands of the task. This change in 1/*f* is consistent with increased inhibition following the onset of a stimulus, and likely reflects a disruption of ongoing excitatory activity proportional to processing demands. Finally, these findings contradict the central assumption of baseline normalization strategies in time-frequency analyses, namely that background EEG activity is stationary across time. As such, they have far-reaching consequences that cut across several subfields of neuroscience.

**Significance statement:** Interest in the functional role of the 1/*f*-like background brain activity has been growing exponentially in neuroscience. Yet, no study to date has demonstrated a clear relationship between information processing and 1/*f* activity by investigating event-related effects on its parameters. Here, we demonstrate for the first time that stimuli induce rotational changes in 1/*f* activity, independent from the occurrence of event-related potentials. Not only do these findings suggest that the assumption of stationary background activity in the analysis of neural oscillations is untenable, but they also suggest the presence of large-scale inhibition following stimulus onset, the magnitude of which is dependent on the demands of the task. These findings have far-reaching consequences that cut across several subfields of neuroscience.

## 1. Introduction

When examined in the frequency domain, the power of neural activity decreases as a function of frequency, following a power law of the form 1/*f*^x^, where *f* denotes frequency and *x* is an exponent that determines the steepness (or slope) of the drop-off in power across frequencies (Pritchard, 1992; He, 2014). This 1/*f*-like shape of the power spectrum reflects the non-oscillatory, or aperiodic, background activity of the brain (Donoghue et al., 2020). This ubiquitous broadband feature was long regarded as noise of no interest that should be corrected for in order to decontaminate the true oscillatory signal of interest (Cohen, 2014; Herrmann et al., 2014; Gyurkovics et al., 2021). However, 1/*f* activity has recently been recognized as a potential source of functionally relevant information in and of itself (He et al., 2010; He, 2014; Gao et al., 2017; Donoghue et al., 2020; Clements et al., 2021). In particular, the steepness (exponent) of 1/*f* activity has been linked to shifts in the balance between excitatory and inhibitory synaptic inputs in local neural circuits, with steeper spectra reflecting a dominance of inhibition over excitation (Gao et al., 2017).

If the shape of 1/*f* activity is indeed a sensitive marker of the level of excitation/inhibition in the cortex, and is therefore related to information processing, the occurrence of a salient stimulus should induce changes in this broadband spectral feature compared to the pre-stimulus state. To the best of our knowledge no study to date has satisfactorily addressed this question directly. While findings from invasive (Miller et al., 2009; Hermes et al., 2015; Podvalny et al., 2015) and non-invasive recordings (Wainio-Theberge et al., 2021) suggest that there might be event-related changes in such activity, these results are difficult to interpret in terms of neurophysiological changes linked to 1/*f* activity because they do not account for the concurrent emergence of event-related potentials (ERPs)^1^.

Salient stimuli and other experimental events reliably elicit ERPs, which are thought to reflect a mixture of aperiodic, transient signals added on top of background activity, and oscillatory activity becoming phase-locked to the event (Makeig et al., 2002; Min et al., 2007; Sauseng et al., 2007; Mishra et al., 2012). As such, this additional activity likely also has a broadband representation in the frequency domain which, when added to ongoing background activity, will confound estimates of 1/*f*-related parameters in the post-stimulus period. Consequently, the frequency content of the ERPs needs to be estimated and removed, to avoid the trivial conclusion that the shape of the power spectrum changes in response to an event simply because the event elicits additional activity. In the present study, we provide an in-depth analysis of stimulus-related changes in 1/*f* activity in two data sets using auditory perception tasks with varying attentional demands. We also illustrate procedures for disentangling genuine event-related changes in background activity *induced* by the event, from ERPs *elicited* by the event. We uncovered a small, but reliable event-related change in 1/*f* activity, consistent with an increased post-stimulus dominance of inhibition. The magnitude of this change also scaled with the attentional demands of the tasks, adding to the growing body of evidence highlighting the functional significance of 1/*f* activity, which was essentially considered noise for decades.

Finally, it is also important to note the methodological implications of event-related changes in 1/*f* activity, which pertain to the analysis of event-related spectral perturbations using time-frequency analysis. To uncover changes often interpreted as oscillatory, this technique requires baseline normalization, which is intended to correct for ongoing oscillatory and non-oscillatory activity pre-stimulus (Gyurkovics et al., 2021). Notably, however, baseline correction methods typically assume that the non-oscillatory part of background activity (1/*f* activity) is stationary across time, and that therefore all baseline corrected post-stimulus changes reflect oscillations or ERPs. Our study constitutes the first empirical investigation of the validity of this assumption. As such, our findings have implications for all subfields of neuroscience that use time-frequency analysis to isolate oscillatory changes.

## 2. Method

### 2.1 Participants

Data collected for a different project (described in Clements et al., 2022) were re-analyzed for the current study. Participants completed two auditory tasks, a passive listening task and an oddball task in separate sessions. Only participants who completed both tasks (n = 23, mean age = 22 ± 2.5, 60.9% female) were analyzed here to make findings across tasks and conditions more comparable. Notably, the results of the passive auditory task do not change when all participants who took part in that wave of data collection (34 young and 12 older adults; n_total_ = 46, mean age = 35 ± 22.2, 56.5% female) are included in the analyses.

### 2.2 Stimuli and Procedure

As mentioned, the 23 participants completed a passive auditory task in which they sat quietly with their eyes open while listening to randomly occurring tone sounds (pips). A 75-ms long, 500 Hz sinusoidal tone was used as the pip, and it was played binaurally from two speakers outside the participant’s field of vision. Participants were told that they would hear pips, but they did not have to respond to them in any form. There were 25 pips in a block, with randomly jittered inter-stimulus intervals (ISIs) between 5 and 10 s. A block lasted approximately 2-3 minutes. Participants had to maintain fixation on a cross presented at the center of a computer monitor throughout the block.

Participants also completed 6 blocks of an auditory oddball task with their eyes open. They heard 25 pips per block (total = 150). However, approximately 20% of the pips (rare pips, on average 30 trials) differed in pitch from the other 80% (frequent pips, on average 120 trials). The pip (450 Hz or 500 Hz) assigned to the rare and frequent group was counterbalanced across participants. All pips were 75 ms in duration. Participants were instructed to mentally count the rare pips and report their number at the end of each block. No other response was required. ISIs were 5-10 s, similar to the passive task.

In the oddball task, trials were sorted into three conditions based on the trial sequence (Clements et al., 2022) to control for well-established sequential effects in oddball paradigms (Brumback et al., 2005). The conditions were: frequent pips preceded by another frequent pip (frequent-<u>frequent</u>, approximately 64% of trials), frequent pips preceded by a rare pip (rare-<u>frequent</u>, approximately 16% of trials), and rare pips preceded by a frequent pip (frequent-<u>rare</u>, approximately 16% of trials). The first pip in each block was discarded from analysis as it had no previous trial type information, and so were rare pips preceded by other rare pips as this trial type occurred extremely infrequently (on approximately 4% of trials).

### 2.3 EEG Recording and Pre-processing

The EEG and electrooculogram (EOG) were recorded from 64 electrodes (60 EEG electrodes placed according to an extended 10-20 system (Jasper, 1958; Acharya et al., 2016), and 4 EOG electrodes placed above and below the right eye, and 2 cm outside the outer canthi of each eye while the participants completed the tasks, using a BrainAmp recording system (BrainVision Products GmbH). EOG and EEG were recorded with a sampling rate of 500 Hz referenced to the left mastoid on-line. The EEG was subsequently re-referenced to the average of the left and right mastoids off-line; vertical and horizontal EOG derivations were obtained by subtracting the below-eye channel from the above-eye channel, and the right-eye channel from the left-eye channel, respectively. Impedance was kept below 10 kΩ during recording. Online, the data were filtered with a 0.5-200 Hz bandpass. Offline, data were pre-processed using the EEGLAB Toolbox (version: 13.6.5, Delorme & Makeig, 2004) and custom MATLAB 2019b scripts (The MathWorks Inc., Natick, MA, USA). A 30 Hz low-pass filter was applied. EEG and EOG data were then segmented into 5120 ms (2560 points) long epochs around pip onset (−2558 ms to 2560 ms, with the onset of the pip being 0 ms). Any EEG channel with variance at least 100 times larger than the median variance of all other EEG channels was considered a bad channel and interpolated as the mean of surrounding good channels. Epochs with amplifier saturation were excluded from further analysis. Vertical (blinks) and horizontal (saccades) ocular artifacts were corrected using the procedure described in Gratton et al. (1983), based on the eye channel recordings. Data from electrodes Fp1 and Fp2 were excluded from analyses as they often contained residual ocular artifacts. Finally, all epochs with voltage fluctuations at any EEG channel exceeding 200 μV after eye movement correction were discarded.

For the frequency analysis of event-related changes in 1/*f* scaling, the pre-event window was defined as the 1024-ms (512 points) long window immediately preceding the onset of the stimulus (−1022 to time point 0), and the post-event window as the 1024-ms (512 points) long window immediately after stimulus onset (2 ms to 1024 ms). The Fourier transform of activity in these time windows was taken on each trial at each electrode for every subject, using MATLAB’s built-in fft() function (Fig. 1). Spectral resolution was 1/1.024 Hz (0.977 Hz). The resulting power spectra were then averaged across trials for the two windows separately within each participant. Frequencies below 1.95 Hz and above 25.39 Hz were removed from further analysis to avoid frequencies whose power estimates were based on fewer than 2 cycles and to ensure that no frequencies affected by the low-pass filter were included, respectively. Then, the resulting spectra were decomposed into oscillatory (periodic) and non-oscillatory (aperiodic) components in semi-log space, using the FOOOF algorithm (v1.0.0, Donoghue et al., 2020)^2^. The aperiodic component of a frequency-domain signal represents 1/*f* activity in the absence of any narrowband oscillations, and the Y intercept (offset) and slope (exponent) of this component were estimated for each spectrum. 1/*f* activity (the aperiodic component) at each electrode for each subject, before and after the event, was then reconstructed in linear space as 10^(ß + *x*log10(*f*))^, where *ß* is the offset in log space, *f* is frequency, and *x* is the slope of the curve. The offset value in linear space (10^ß^), the slope (with a negative sign, i.e., *x*), and the reconstructed aperiodic component in each time window, at each electrode for every participant were retained for further analyses.

**Figure 1.**
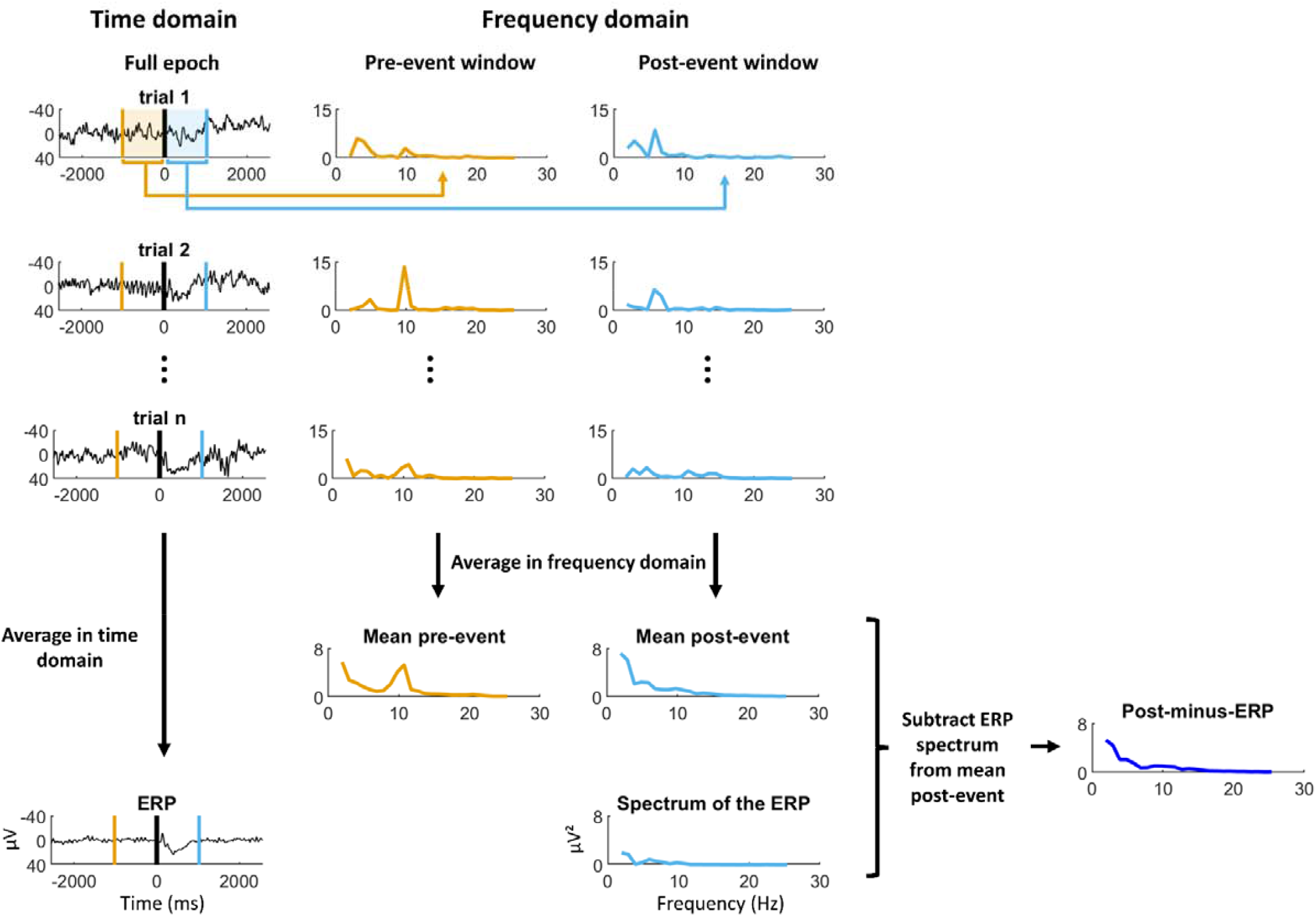
An illustration of the various steps during data preparation for a single subject, in a single condition, at a single electrode. For illustration purposes, the frequent-<u>rare</u> condition was selected at electrode Pz, although this process was completed at all electrodes. The mean pre-event, mean post-event, and post-minus-ERP spectra were then decomposed into aperiodic and periodic components using FOOOF. Negative is plotted up in the time domain. ERP = event-related potential

To account for the presence of ERPs in the post-event period, the spectrum of the ERPs was also quantified for each condition within each participant using the fft() function, i.e., the power spectra of the cross-trial time-domain averages were calculated, instead of calculating average power spectra across trials. These spectra were then subtracted from the post-event spectra to yield post-minus-ERP difference spectra (Fig. 1). This is conceptually identical to subtracting the ERPs from single-trial activity in the time domain. Notably, these post-minus-ERP difference spectra were subsequently also deconstructed by FOOOF, and their offset and exponent values saved. The reconstructed aperiodic components of the pre-event spectra were then subtracted from the reconstructed aperiodic components of the difference spectra to yield an estimate of the event-related changes in aperiodic activity (1/*f* shift) unconfounded by the presence of ERPs. The parameters of this event-related aperiodic change estimate – quantified as the difference in exponent and offset between the two aperiodic components described above - were then contrasted across conditions within the oddball data set to investigate the attention-dependence of the shift in background activity.

### 2.4 Statistical Analysis

All statistical analyses were based on non-parametric methods, using a truncated permutation distribution approach. First, we tested whether there was an overall significant difference, across all electrodes, between the pre- and post-event offset and slope estimates of the 1/*f* component of the spectrum, as computed using FOOOF. To this end, the grand mean post-minus-pre difference value across all electrodes and all participants was compared against a null distribution of difference values based on 10,000 permutations, separately for each parameter (offset and slope). On each permutation, the sign of the mean difference for a given parameter across electrodes was flipped for a random half of the subjects, and the grand mean difference was calculated on this new sample. This resulted in a null distribution with 10,000 grand mean values, in which the 2.5^th^ and 97.5^th^ percentiles were identified. If the observed grand mean difference value fell outside of the range between these two percentiles, it was considered significant at *p* < .05 (two-tailed). For the oddball paradigm, this procedure was run separately for each condition (frequent-<u>rare</u>, rare-<u>frequent</u>, frequent-<u>frequent</u>) for these initial analyses.

Second, the pre-event offset and slope values were contrasted with the offset and slope values of the post-minus-ERP difference spectra, using the same permutation testing-based approach as above, to examine if there were significant event-related changes in aperiodic activity even after accounting for ERPs.

Next, three progressively more complex models of post-event spectral activity were contrasted to establish whether accounting for post-event changes in 1/*f* activity explained a significant amount of variance above and beyond the contribution of the ERPs. In the first model, post-event power was predicted from pre-event power at each frequency within each participant.

**Model 1**: post-event spectrum = pre-event spectrum

In the second model, the spectrum of the ERPs was also added as a predictor.

**Model 2**: post-event spectrum = pre-event spectrum + ERP spectrum

In the final model, post-event power was predicted as a sum of pre-event power, the spectrum of the ERPs, and a shift in 1/*f* activity, estimated as the difference between the FOOOF-reconstructed 1/*f* components of the post-minus-ERP difference spectrum and the pre-event spectrum.

**Model 3**: post-event spectrum = pre-event spectrum + ERP spectrum + 1/*f* shift

In all models, the predictor(s) was (were) simply subtracted from the post-event spectrum to obtain residuals instead of using a regression approach (i.e., there was no fitting of weights given to each component, but instead fixed weights of 1 were assigned to all components). Squared residuals were then summed across electrodes, participants, and frequencies below 7 Hz and above 13 Hz, to eliminate the alpha range (greyed out in all figures). In the alpha frequency band, a predicted suppression of activity occurred post-event which was not accounted for in any of our models as it was of no relevance to our research question. These resultant models were then compared by calculating *F*-ratios in a stepwise fashion (Model 1 vs. Model 2, and Model 2 vs. Model 3) with degrees of freedom *n*-2, *n*-3, and *n*-5 for Models 1, 2, and 3, respectively, where *n* is equal to the sample size. These values correspond to *n* minus the number of predictors in each model minus 1, accounting for the DC offset (time domain mean) which is estimated during the Fourier transform. One additional degree of freedom was expended in model 3, as the 1/*f* components of each spectrum were reconstructed based on 2 parameters (offset and slope) that were estimated from the data. This represents a conservative approach to the calculation of degrees of freedom given that models 1 and 2 did not technically involve the estimation of any parameters as no fitting was performed to obtain residuals. Critical *F*-values were calculated at α = 0.05 for each model comparison, and any *F*-ratios exceeding these critical values were considered to indicate a significant reduction in error variance across models.

Finally, to compare the magnitude of the estimated 1/*f* change across oddball conditions, the difference between pre-event offset and slope values and the offset and slope values of the post-minus-ERP difference spectra were calculated for each condition (quantifying genuine 1/*f* change after accounting for ERPs). Then these difference values were compared between the frequent-<u>frequent</u> and rare-<u>frequent</u>, and the frequent-<u>frequent</u> and frequent-<u>rare</u> conditions, using the same permutation testing-based approach as above. The α level was adjusted using the Bonferroni approach to account for the two pairwise comparisons. Importantly, to equalize the number of trials across conditions, observations were randomly selected from each condition to match the lowest trial number across all 3 conditions within each subject. This random selection with replacement was completed 100 times, and the mean spectra across these bootstrapped samples was only used in the cross-conditional comparisons described here.

The code used to run the preprocessing steps and the analysis pipeline is available at https://osf.io/nq8jk/.

## 3. Results

### 3.1. Event-Related Rotational Shift in Aperiodic Background Activity Above and Beyond the Contribution of ERPs

Fig. 2A shows the pre- and post-event spectra across conditions. Differences are present in the alpha range (corresponding to alpha suppression, greyed out because it is not relevant to the central question of the current paper), and at frequencies below alpha. These latter differences that are not confined to a narrow band can be summarized using the best fit aperiodic components from FOOOF (dashed lines). Permutation-based analyses confirmed that both aperiodic slope and offset values were significantly different at *p* < .05 between the two time-windows, in all conditions. However, this difference in aperiodic shape parameters could be due to the presence of ERP activity in the post-stimulus spectrum, which is likely to be particularly prominent at low frequencies. Therefore, we subtracted the spectra of the corresponding ERPs from the post-event spectra in each condition and re-computed the FOOOF parameters on the residual spectra. The slope values of these residual post-event spectra (post-minus-ERP) were still found to be significantly different from the slope values of the pre-event spectra in every condition (Fig. 2B). This change in slope is consistent with a clockwise rotational change in the shape of the spectrum (Donoghue et al., 2020) from pre- to post-event. In contrast, after subtracting the ERP spectrum from the post-stimulus spectra, differences in the offset parameter between the pre- and post-stimulus spectra did not reach significance.

**Figure 2.**
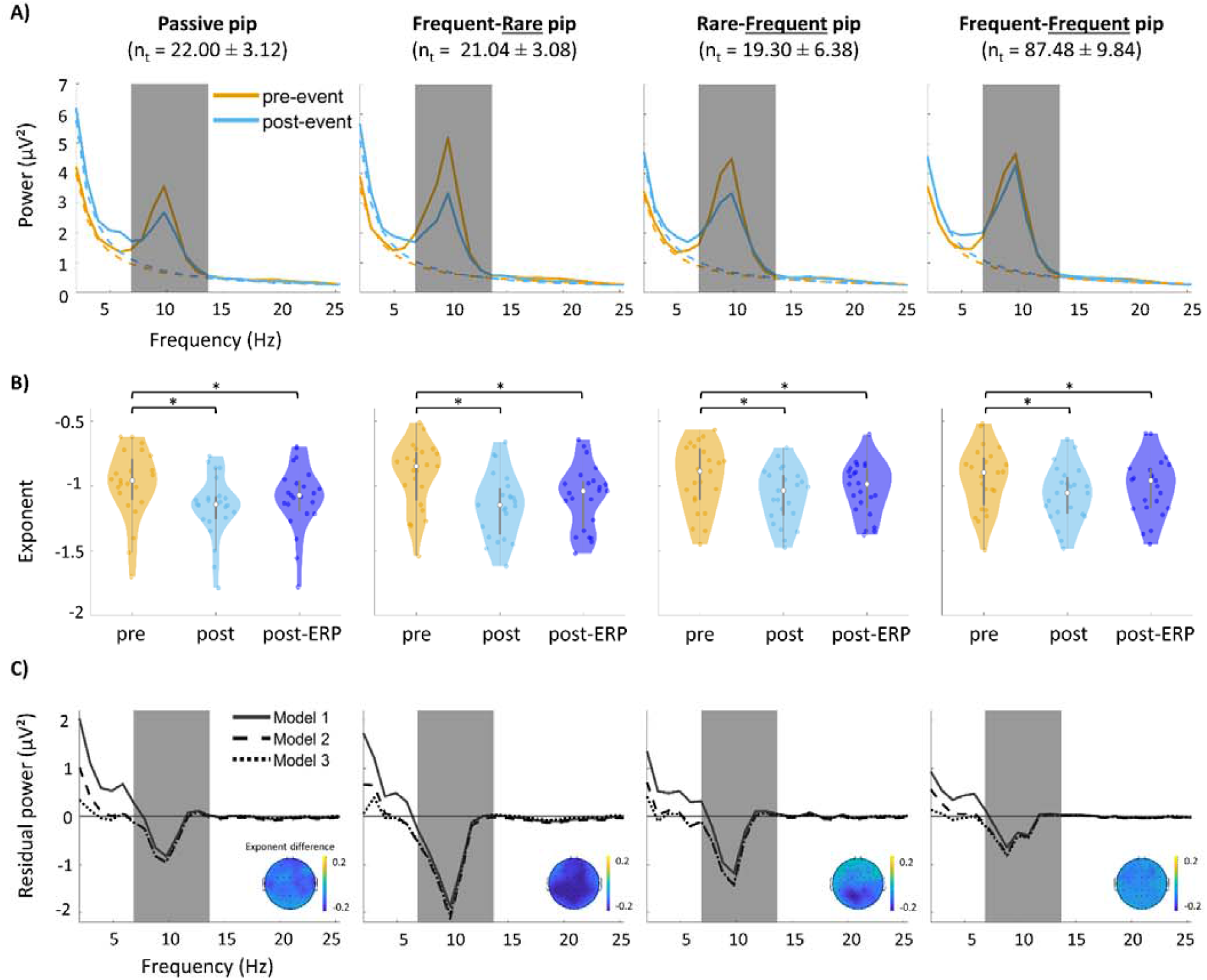
Post-event changes in spectral content across conditions. The mean trial number (± SD) in different conditions is indicated in parentheses below the condition labels. A) Mean pre- and post-event power spectra in different conditions (solid lines), along with their best aperiodic fit from FOOOF (dashed lines). Greyed out frequencies (7-13 Hz) indicate the alpha range which was of no relevance to our research question of interest, and was excluded from formal calculations of model fit (see main text). B) The distribution of exponent values across participants in different time windows across conditions. „Pre” indicates the exponent of the pre-event spectrum, „post” indicates the exponent of the post-event spectrum, and „post-ERP” (post-minus-ERP) indicates the exponent of the post-event spectrum following the subtraction of the ERPs. Asterisks indicate comparisons significant at p < .05 based on permutation testing. C) Residual post-event power after accounting for different sets of predictors. Predictors were pre-event power (Model1, solid line), pre-event power and ERP spectrum (Model 2, dashed line), and pre-event power, ERP spectrum, and estimated 1/*f* shift (Model 3, dotted line, see main text). Greyed out frequencies (7-13 Hz) indicate the alpha range. Insets indicate the scalp distribution of the exponent difference between pre-event power and post-event power minus the contribution of ERPs, i.e., the exponent shift in 1/*f* activity not accounted for by ERPs.

The impact of this stepwise subtraction of the pre-event spectra, the ERP spectra, and then finally of the 1/*f* components can be seen in Fig. 2C. The traces represent the residual variances in post-event power after removing the contribution of these different sets of predictors in a stepwise fashion, corresponding to the residuals of Models 1, 2, and 3 introduced above in section 2.4. The black trace corresponds to the residual spectrum obtained with Model 1, revealing substantial unexplained power at low frequencies (as well as in the alpha range, which is consistent with suppression following pip onset.) Residuals at low frequencies are greatly reduced with Model 2 (dashed line) compared to Model 1; however, they still clearly deviate from 0. This is consistent with an actual shift in 1/*f* activity from pre- to post-event. To illustrate this point, the dotted trace in Fig. 2C shows the residual post-stimulus spectrum obtained with Model 3. Compared to Model 1 and 2, application of Model 3 results in decreased unexplained variance in power at low frequencies, and often at higher frequencies as well. It is likely that any remaining residual variance could be attributed to cumulative noise in the estimation of both the ERPs and the aperiodic components.

Formal analyses of the change in error variance outside the alpha rage across models are presented in Table 1. These analyses of variance confirmed that unexplained (i.e., error) variance in the post-stimulus spectrum significantly decreased after the contribution of the ERP spectra was accounted for, and, crucially for the purposes of this paper, decreased further after accounting for a change in 1/*f* activity.

**Table 1.**
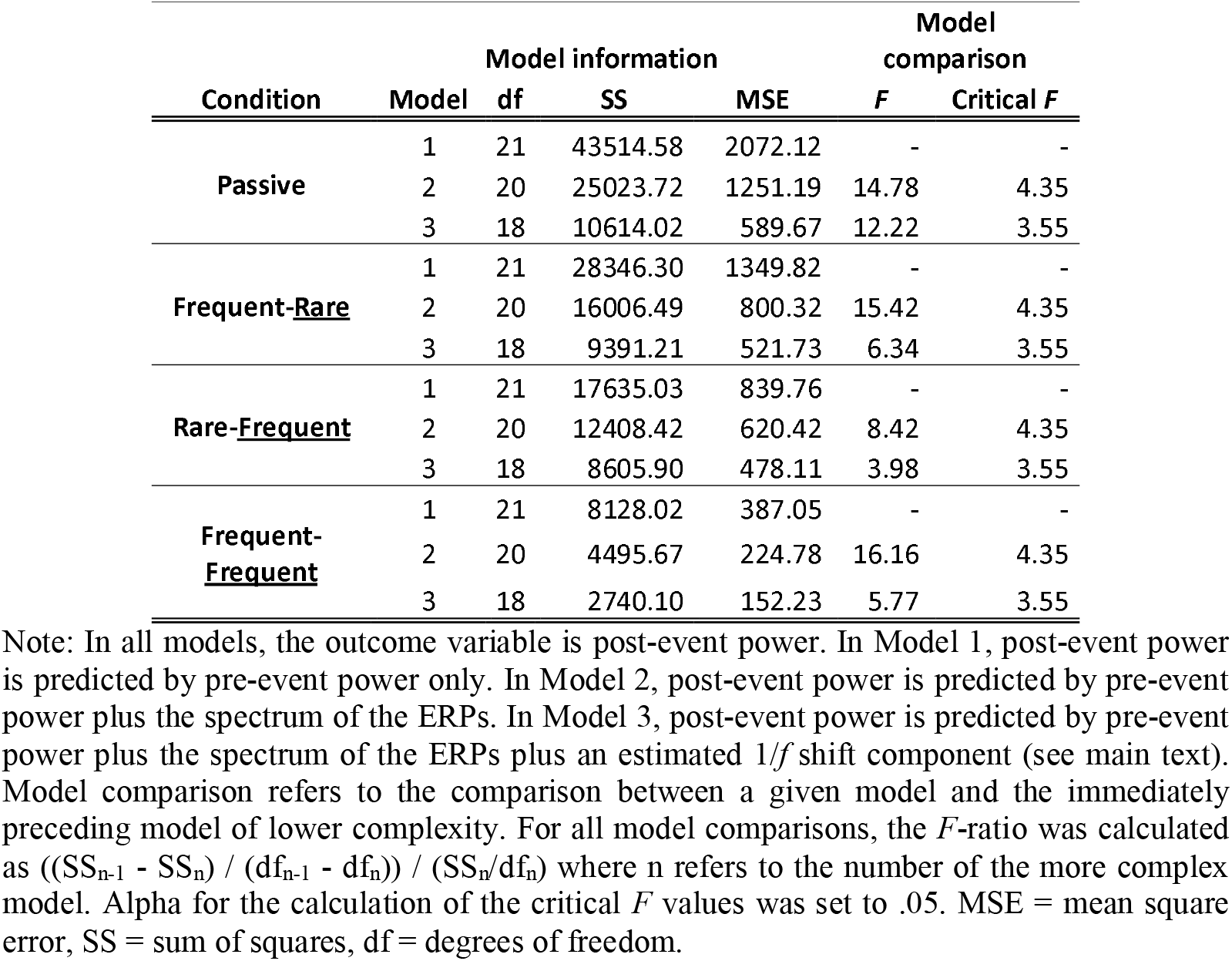
Predictive models of post-event power across conditions.

Note that the alpha range was excluded from the analyses of variance because we did not expect post-event alpha activity to be accounted for by any of our models, as alpha suppression reflects a modulation of the amplitude of ongoing alpha oscillations that is not part of the ERP or of non-oscillatory background activity. Accordingly, accounting for the ERP did not decrease the difference between pre- and post-event activity in the alpha range (Model 1 vs Model 2 in Fig. 2C). Notably, however, changes following ERP removal were pronounced in the theta range, suggesting that a substantial portion of event-related theta activity is elicited by the event, and does not reflect the modulation of ongoing oscillatory activity.

Taken together, these findings suggest that there is a small but reliable increase in the steepness of aperiodic background activity in response to simple auditory stimulation, and that this clockwise rotational shift in the spectrum is not explained by activity elicited by the stimuli (i.e., by ERP activity).

The analyses presented above were based on averaging spectra across all electrode locations. When individual electrodes are considered separately, the scalp distributions of the changes in the slope parameter after accounting for ERPs (insets in Fig. 2C) indicate that these changes (pointing at a rotational shift in the aperiodic EEG spectrum) are broadly distributed in most conditions, suggesting global rather than localized effects.

### 3.2. The Magnitude of the Rotational Shift in 1/f Activity is Attention-Dependent

We further compared the magnitude of this rotational shift across conditions in the oddball task data (Fig. 3). Two conditions require the reconfiguration of ongoing processing because of a change in the stimulus that was presented, rare-<u>frequent</u> and frequent-<u>rare</u> trials. These conditions were each contrasted against the frequent-<u>frequent</u> trials, which require no such change in processing. These two conditions differ, however, on whether the change was relevant to the subject’s task. Differences in exponent (i.e., slope) values between the post-minus-ERP spectra and the pre-event spectra (an estimate of the 1/*f* rotational shift) in the two stimulus-change conditions were compared with the same 1/*f* rotational shift estimate on the frequent-<u>frequent</u> trials in a pairwise fashion, using permutation testing (Fig. 3A, B). The contrast between frequent-<u>rare</u> (where stimuli are different, task relevant, and have low global probability) and frequent-<u>frequent</u> trials (which do not entail any of these three features), reached significance at *p* < .025, indicating that the increase in steepness was larger after rare trials preceded by a frequent trial than after frequent trials preceded by a frequent trial, with a widespread scalp distribution (Fig. 3C). The contrast between frequent-<u>frequent</u> and rare-<u>frequent</u> trials did not reach significance. These cross-conditional differences suggest that the change in 1/*f* activity reflects functionally relevant processes sensitive to a combination of stimulus change and task relevance of the stimuli.

**Figure 3.**
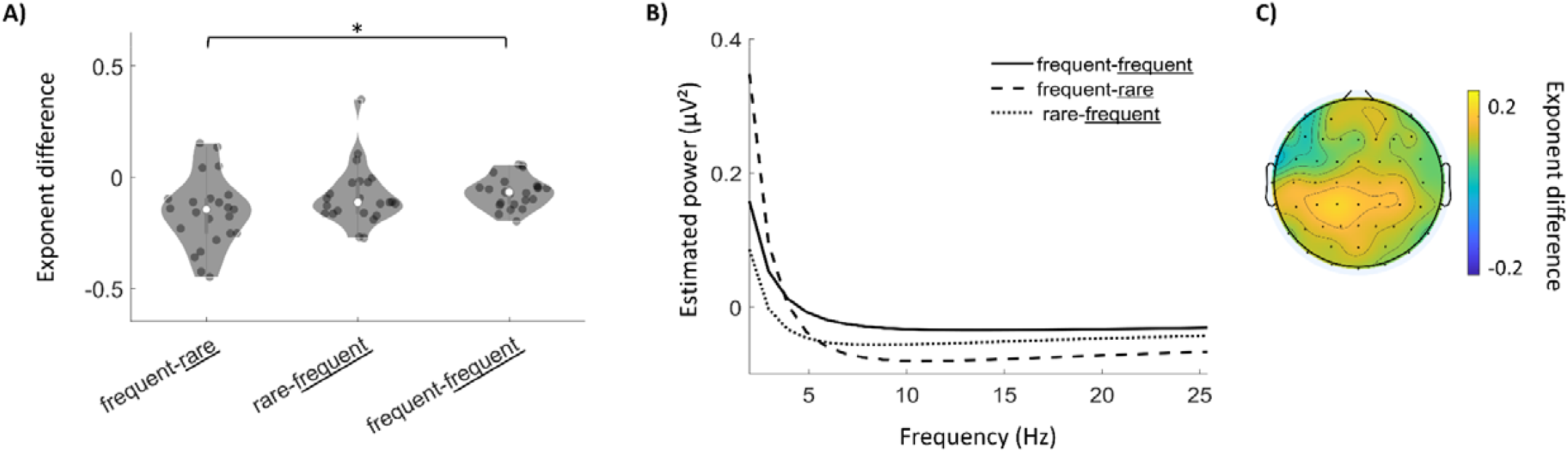
Conditional differences in the magnitude of the post-event rotational shift in 1/*f* activity in the oddball data set. A) The distribution of the difference between the post-minus-ERP exponent and the pre-event exponent across participants, for the three oddball conditions. B) The lines represent the difference between estimated pre-event 1/*f* activity (aperiodic component) and estimated post-event 1/*f* activity after accounting for the contribution of ERPs (post-minus-ERP). Negative values occur because the curves are the result of subtractions. Trial numbers were equated for the calculation of these slopes. For the scalp distribution in the inset, the difference between the exponent of pre-event power and the exponent of post-minus-ERP power (i.e., an estimate of genuine 1/*f* change) in the frequent-frequent condition (rightmost violin in A) were subtracted from the same difference values in the frequent-rare condition (leftmost violin in A).

## 4. Discussion

In the present study, we investigated event-related changes in the 1/*f*-like non-oscillatory background activity of the brain during two simple auditory tasks. We found a set of complex changes in the spectral composition of neural activity in response to auditory stimuli in our datasets. Importantly, in addition to an expected post-event decrease in oscillatory activity in the alpha range (i.e., alpha suppression e.g., Klimesch, 1999; Klimesch et al., 2007; Palva and Palva, 2007; Foxe and Snyder, 2011; Feng et al., 2017), changes in both the offset and the slope of 1/*f* background activity (as estimated using the FOOOF algorithm; Donoghue et al., 2020) were also consistently observed across tasks. Changes in the slope parameter were still detectable, although substantially reduced in magnitude after removing the frequency-domain contribution of ERPs, reflective of the transient activity elicited by the event. This supports the presence of genuine alterations in the ongoing non-oscillatory activity of the brain following stimulation, The observed spectral shift was largest when participants encountered an oddball stimulus (i.e., a stimulus that is both rare and task relevant), suggesting that the change in 1/*f* activity is associated with attentional processes.

We found that the steepness of 1/*f* activity increased after auditory stimulation, resulting in a clockwise rotational shift of the spectrum which was characterized most prominently as an increase in low frequency activity following stimulus onset. Within recent frameworks that link the exponent of non-oscillatory activity to an excitatory-inhibitory balance, this rotational change is consistent with a shift towards a more inhibitory regime (Gao et al., 2017; Waschke et al., 2021). This increased inhibition was widespread, and not confined to a focal scalp distribution across conditions (see inset scalp topographies in Fig. 2C). It is possible that this inhibitory dominance reflects the disruption of ongoing activity, as part of the brain’s response to the occurrence of a new stimulus. Overriding existing representations and establishing new ones have been linked to brief occurrences of lower frequency activity (Gratton, 2018). When considering the nature of the activity contributing to the rotational shift, it is worth noting that the changes observed in our data set are unlikely to reflect stimulus-induced modulations of *oscillatory* activity in particular frequencies simultaneously, as they were not constrained to narrowband peaks within the spectrum. Indeed, narrowband peaks were largely only observed in the alpha range. Similarly, the observed changes cannot be taken to reflect stimulus-*elicited* activity (Cohen, 2014), as they did not contribute to the ERP. As such, we conclude that our findings reflect stimulus-*induced* changes in already ongoing *non-oscillatory* background activity.

Our findings also suggest that this stimulus-induced shift in 1/*f* activity was largest when participants encountered a rare oddball stimulus immediately after a frequent standard stimulus. This is in line with the idea that the rotational shift reflects stimulus-induced inhibition that helps override pre-existing representations, because this process should be the most pronounced when a rare stimulus (oddball tone) is encountered, the representation of which differs from the established, dominant representations (frequent tones). The rotational shift was not dependent on whether a frequent tone was preceded by a rare tone or another frequent tone, possibly because re-establishing the dominant representation caused less disruption to ongoing processing. Taken all this into account, we propose that the shift toward slower activity observed in these data reflects a global, non-focal inhibitory mechanism that temporarily halts ongoing processing to allow for the subsequent clearance of outdated representations and the establishment of new ones, thus contributing to a broader orienting response (Sokolov, 1990). This notion is similar to the transient, global blocking of all motor output immediately after a novel stimulus, which is posited to occur by various models of attentional control (Munakata et al., 2011; Wiecki and Frank, 2013; Wessel and Aron, 2017; von Bastian et al., 2020) although the mechanism proposed herein is more general, as it is not limited to motor outputs. Indeed, our task required no motor responses, and the scalp topography of the 1/*f* shift does not indicate an origin in motor-related areas.

The post-stimulus suppression of alpha activity, which was observed to be clearly oscillatory both pre- and post-event in our study, is also thought to index the destabilization of current representations to allow for the instantiation of new ones (Gratton, 2018; Clements et al., 2022). However, the selective attention mechanisms likely reflected by alpha activity are more local and specific to goal-relevant areas than the large-scale and seemingly indiscriminate global inhibition suggested by the scalp distribution of the 1/*f* shift. As such, the former is assumed to be more closely tied to top-down, goal-related control of attention than the latter.

Notably, a recent study by Waschke et al. (2021) found an attention-dependent decrease, as opposed to increase in the steepness of 1/*f* activity in a multimodal change detection task in scalp-EEG recordings. Those findings, however, did not reflect stimulus-induced changes, as the spectra were quantified in time windows far removed from trial onset, during sustained attention to continuous auditory or visual stimulation. It is possible, and indeed likely, that the momentary inhibitory shift induced by an event, as observed in our study, is then overtaken by a dominance in excitation if the stimulation persists, especially in areas contributing to the processing of information in the target modality (Harris and Thiele, 2011), as observed by Waschke et al. (2021). Consequently, their findings are complementary to the results presented here, and suggest that the re-orienting of attention to new stimuli and the sustained attention within a modality engender changes in the opposite direction in ongoing brain activity.

Our findings also have profound implications for the analysis of task-based neural data in the time-frequency domain. In these analyses, baseline normalization is applied to emphasize event-related changes in power at various frequencies (Cohen, 2014; Herrmann et al., 2014; Gyurkovics et al., 2020). Normalization strategies typically involve subtracting from or dividing power values in an epoch by an estimate of baseline activity, most frequently based on the pre-event period. These strategies assume that 1/*f* activity does not change from the baseline pre-event period to the post-event period. As such, removing an estimate of the former from the latter is assumed to recover event-related changes in oscillatory activity occurring against the backdrop of consistent background noise. The current results, however, run counter to this assumption, and highlight that pre-event and post-event activity may differ in broadband characteristics, even after accounting for ERPs. As such, researchers must be cautious in interpreting significant changes compared to baseline as oscillatory changes in time-frequency analyses, especially at low (below alpha) frequencies because they may simply reflect a shift in 1/*f*-like background activity that is non-oscillatory in nature, or may be confounded by it. The analytic pipeline outlined in this paper provides a means to examine if such a shift is present in a given data set, which could potentially compromise the interpretability of low frequency findings in time-frequency analyses. However, note that our analyses are most appropriate for paradigms with ITIs long enough (∼1024 ms or preferably more) to provide good frequency resolution both before and after a target stimulus.

### 4.1 Conclusions

In the present study, we found that the steepness of 1/*f*-like non-oscillatory background activity in the brain increases after auditory stimulation, and this broadband change is separable from the phenomenologically similar contribution of ERPs. The observed rotational shift in the power spectrum is consistent with a global increase in inhibitory activity following a rare, task-relevant stimulus, and likely represents the momentary disruption of ongoing processing to enable further stimulus-specific processing. Infrequent stimuli induced a larger broadband shift than frequent stimuli, underlining the potential functional significance of the observed spectral change. Our findings also suggest that baseline normalization strategies common in time-frequency analysis incorrectly assume that background activity is stationary across task-based epochs, and this may render findings difficult to interpret.

## Footnotes

1 Notably, Hermes et al. (2015) did observe a broadband shift at high frequencies (50-200 Hz) in two subjects following visual stimulation using ECoG after regressing out ERPs, but the lower end of the spectrum which has much higher signal-to-noise ratio in noninvasive scalp recordings compared to higher frequencies and is likely to contain most of the power of the ERPs, was not examined.

2 The following settings were used for spectral parametrization: peak_width_limits=[2,8], max_n_peaks=1, min_peak_height=0.3, peak_threshold=2.0, aperiodic_mode=‘fixed’. The maximum possible number of extracted peaks was set to 1 because the algorithm rarely selected more peaks when less restrictive settings were used, and we aimed to avoid increasing variability across participants by allowing noisy peaks to be selected for some participants but not for others. Importantly, however, our results do not change if the max_n_peaks parameter is set to 2, 3, or higher numbers.

